# “Role of Tad Pili during the transition from Planktonic to Biofilm State in *Bradyrhizobium diazoefficiens* USDA 110”

**DOI:** 10.1101/2025.01.14.633045

**Authors:** J. Iglesias, D. Colla, JS. Serrangeli, M. Lozano, O. Falduti, D. Brignoli, I. Medici, MJ. Al-thabegoithi, AR. Lodeiro, PL. Abdian, N. Paczia, A. Becker, A. Soler-Bistue, J. Perez-Gimenez, EJ Mongiardini

**Author notes:** Iglesias J. and Colla D. contributed equally to this work. Present address: Fundación Instituto Leloir, Ciudad Autónoma de Buenos Aires, Argentina (JS. Serrangeli).

## Abstract

Free-living soil bacteria may exist in two states: planktonic as swimmer cells, and sessile within biofilms. In biofilms, bacterial cells are embedded into an extracellular matrix, which confers protection against environmental stresses and extends bacterial survival. The transition from planktonic to biofilm state involves surface sensing and attachment mediated by flagella and pili. Here, we explored the role of Tad pili (Type IVc) in surface sensing, adhesion, and biofilm formation in *Bradyrhizobium diazoefficiens*, a nitrogen-fixing symbiont of soybean. Bioinformatic analyses revealed that Tad pili are widely distributed and highly conserved in *Bradyrhizobium*. In other model bacteria, pili deletion leads to decrease in biofilm-forming capacity, but other functions are also affected depending on the microorganism. Surprisingly, deletion of the most conserved genomic cluster encoding Tad pili in *B. diazoefficiens* caused an increased adhesion to abiotic surfaces and loss of movement, indicating a physiological shift towards a biofilm state. These observations suggest a sensor or regulatory role for this apparatus that could affect cell-cell interactions or interactions with the extracellular matrix. Additionally, a connection between pili and c-di-GMP levels in the cell was established. These findings highlight the critical role of Tad pili in *B. diazoefficiens* physiology, providing insights into bacterial adaptability and potential applications in agriculture and biotechnology. Understanding these mechanisms is key to improving biofilm management strategies and generating novel approaches to enhance bacterial viability in soil and inoculants as well as to optimize the symbiosis.

**Importance:** Biofilm formation is critical for bacterial survival in the soil. In this study we analyzed the role of Tad pili in *Bradyrhizobium diazoefficiens* biofilm forming capacity and its connection with the second messenger c-di-GMP, which is involved in regulating the transition between planktonic and sessile states. Since bacteria applied in inoculants are in the planktonic state, whereas bacterial survival is optimal in the sessile (biofilm) state, our findings may contribute to im-prove the switch towards the sessile state in novel inoculant formulations, enhancing bacterial persistence in formulations and soil.

## Introduction

Free-living soil bacteria exist in two main states: planktonic and sessile. Planktonic bacteria are characterized by individual cells that can swim without restriction. By contrast, sessile bacteria often are grouped in biofilms, a community in which bacteria first colonize a surface, produce an extracellular matrix and develop multicellular characteristics, creating a highly organized and resilient structure (1–3). In biofilms, bacterial cells are embedded within a polymeric matrix that may consist of polysaccharides, proteins, and/or nucleic acids, which confer protection against environmental stresses and contribute to cell survival (4).

The ability of bacteria to sense a surface and attach to it is a crucial step during the transition from planktonic to biofilm state. These processes are commonly mediated by bacterial surface appendages, such as flagella and pili, which play critical roles in initiating the formation of bio-films (5, 6). Flagella are primarily involved in motility, allowing bacteria to navigate toward favorable environments, while pili are elongated, filamentous appendages that facilitate different functions (7, 8).

Pili, also known as fimbriae, are dynamic fibers composed of pilin subunits that polymerize and depolymerize, allowing the filament to extend and retract. This dynamic nature enables bacte-ria to probe their environment, attach to surfaces, and form cohesive communities (9, 10). Several types of pili have been identified in bacteria, each associated with diverse cellular functions (11, 12). The versatility of pili underscores their importance in bacterial survival and interaction with other organisms.

The classification of pili encompasses several families categorized according to their structure and function. Among these, Type IV pili (T4P) are the most abundant and diverse in the micro-bial world, present in both Eubacteria and Archaea (11). T4P are involved in a wide range of functions, including twitching motility, DNA uptake during transformation, and virulence in pathogenic bacteria (9, 13, 14). Within this group, the Type IVc subclass (T4cP) is particularly noteworthy. T4cP, also known as Tad pili, are widely distributed among bacteria and are implicated in surface sensing, adhesion, and biofilm formation (15–17).

T4cP are encoded in genomic islands and their pilin subunits typically have low molecular mass (15). They were first characterized in *Aggregatibacter actinomycetemcomitans* as being responsible for tight <u>ad</u>herence to surfaces, hence their name, Tad pili (18). This tight adherence facilitates the formation of robust biofilms, which are critical for the ability of these bacteria to colonize and persist in the oral cavity, contributing to periodontal disease (19). At the same time, T4cP were described in *Caulobacter crescentus* as an apparatus connected to cell cycle progression, highlighting their multifunctional role in bacterial physiology (20).

More recently, Tad pili have been proposed as the main system involved in surface sensing in *C. crescentus*. These pili, together with the flagellum, initiate a cascade of signaling events upon surface contact, which is crucial for developmental transition of bacteria into the biofilm mode of life (21–25). In other bacteria, such as *Agrobacterium tumefaciens*, Tad pili play a pivotal role in adhesion to abiotic surfaces and biofilm formation, which are essential for the bacterial ability to infect plant hosts (26). Similarly, in *Sinorhizobium meliloti* and *B. diazoefficiens* Tad pili are implicated in the initial stages of interaction with plant roots during the establishment of symbiosis, influencing the bacterial competitiveness to form nitrogen-fixing nodules on legumes roots (27–29).

Another key player in the transition mechanism between the planktonic and biofilm states is the second messenger c-di-GMP, a molecule that integrates many stimuli sensed by bacterial receptors (30). This molecule increases its concentration when the bacterium senses a surface, initiating the cellular program that leads to biofilm formation. The synthesis of c-di-GMP is catalyzed by diguanylate cyclases connected to flagella and pili functioning. A mechanical disruption of their function by the presence of a surface act as trigger for the second messenger synthesis. (21, 22).

In the genus *Bradyrhizobium*, which comprises nitrogen-fixing bacteria that form symbiotic nodules in legume plant roots, the presence and role of pili have been previously noted (31). *B. diazoefficiens* has been characterized as possessing adhesive fimbriae; however, neither its physiological role has been explored yet, nor the specific structural gene encoding the 20 kDa pilin subunit has been identified (31). In a previous study, we identified a *tadG* homolog in *B. diazoefficiens* through its similarity to the BJ3) lectin, which might be related to Tad pili (29). This protein is part of a small cluster of genes with high homology to accessory proteins of pseudopilin subunits. Nevertheless, its role in the structure and function of Tad pili was not confirmed.

Here, we present a phylogenetic study of the Tad pili apparatus in the *Bradyrhizobium* genus, focusing on its function in the regulation of the transition from planktonic to sessile state. By characterizing the genetic and structural components of Tad pili in *B. diazoefficiens*, we aimed to elucidate their role in surface sensing, adhesion, and biofilm formation. Also, we report a connection between the c-di-GMP level and the Tad pili in these bacteria. Understanding the regulation of adhesion mechanisms is critical not only for advancing our knowledge of bacterial physiology but also for exploring potential applications in agriculture and biotechnology, where the manipulation of this process and biofilm formation could have significant implications.

## Results

### 1. Tad pilus is widely present and highly conserved in the *Bradyrhizobium* genus

The Tad pilus, or T4cP (21), is widely distributed within *Bradyrhizobium*. It was recently reported that *B. diazoefficiens* USDA 110 exhibits four Tad pili clusters (28), which we renamed from 1 to 4 in accordance with their ordered location on the chromosome. Here, we aimed at the bioinformatic and molecular characterization of these clusters in *Bradyrhizobium. B. diazoeffi-ciens* USDA 110 has two complete and two partial pili clusters. For clarity, all genes within each cluster were named using the nomenclature from *Agrobacterium tumefaciens* C58*, Caulobacter crescentus* N1000*, and Actinobacillus actinomycetemcomitans* D7S-1, and their predicted functions were assigned based on sequence similarity by bi-directional Basic Local Alignment Tool-Protein (BLASTp) comparison (Fig. 1; Supplementary Table S1). When several paralogs were found, the cluster number was added to the names as suffix.

**Figure 1.**
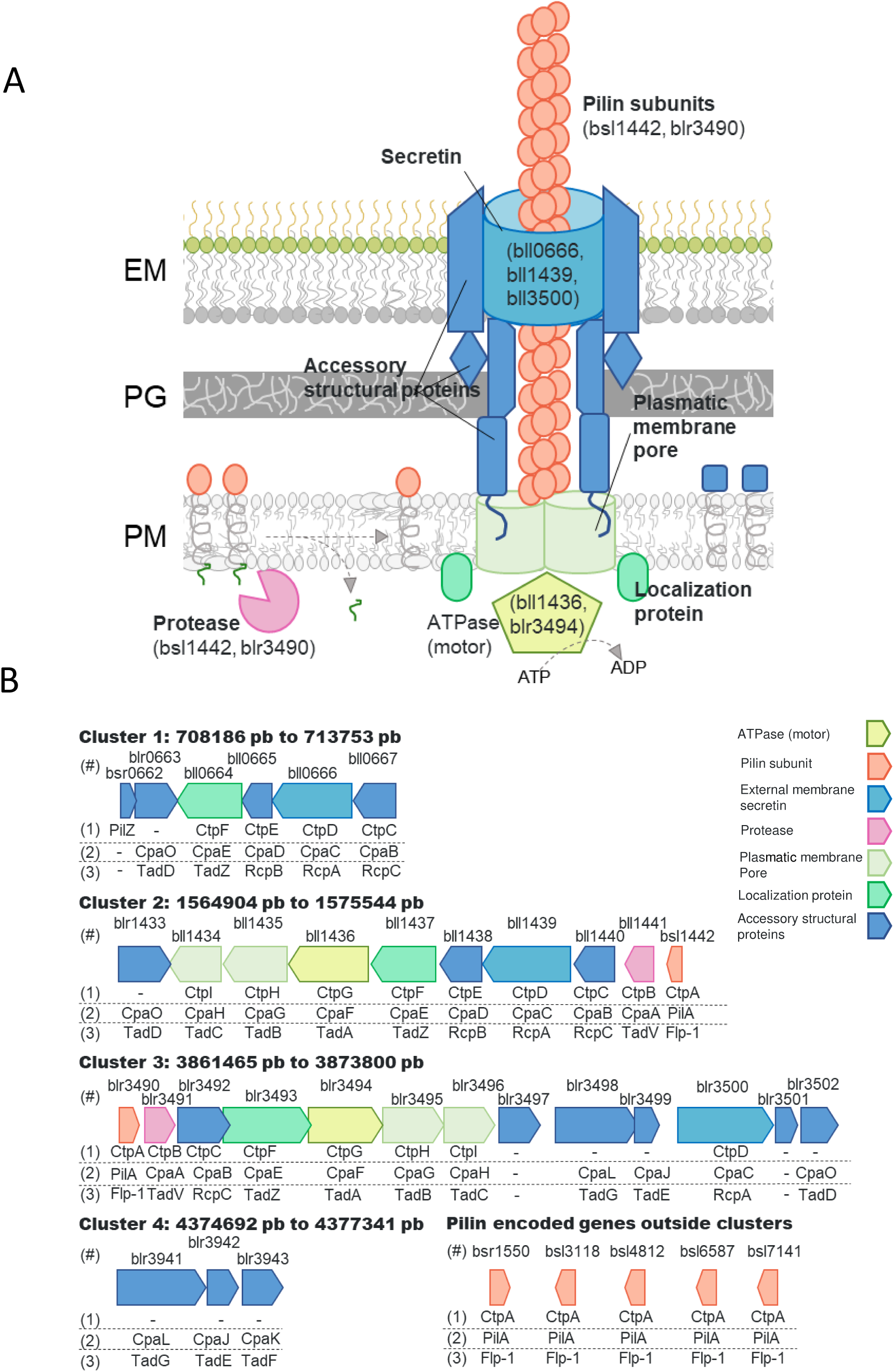
Tad pili clusters encoded in *Bradyrhizobium diazoefficiens* USDA 110. **A.** Scheme of a Tad pilus. **B.** Scheme of the gene clusters related to Tad pili systems in *B. diazoefficiens* USDA 110. The genes assigned to each cluster are indicated below the gene, according to the results shown in Table S1, and using the same nomenclature as in three different bacterial models: (1) *Agrobacterium tumefaciens* C58, (2) *Caulobacter crescentus* N1000, and (3) *Actinobacillus actinomycetemcomitans* D7S−1. Each gene is identified by the locus tag (above each gene box).

A global comparison of the four clusters encoded in *B. diazoefficiens* USDA 110 and the reference genomes used in the function assignment was run using Clinker (Supplementary Fig. S1). Within *B. diazoefficiens* USDA 110, we noticed high similarity between cluster 2 and the partially encoded cluster 1, suggesting its origin by a duplication event. Additionally, cluster 2 shows high conservation in sequence and cluster organization with *A. tumefaciens*, *S. meliloti*, and *C. crescentus*. On the other hand, clusters 3 and 4 display relatively low conservation in comparison to the others.

Furthermore, all genome sequences belonging to the *Bradyrhizobium* genus available in the NCBI database (1505 species, accessed 30/09/2024) (Supplementary Table S2) exhibit multiple, complete or partial gene clusters encoding Tad pili apparatus (Supplementary Fig. S2), whereas 95% genomes have at least cluster 1 and/or cluster 2. The remaining genomes (5%) where we were unable to detect the presence of Tad pili correspond to genomic data at the assembly level of contigs or scaffolds which may be the cause of failed detection. This prevalence suggests a significant role for the Tad pilus in the lifestyle of this rhizobacterium; however, its function has not yet been thoroughly studied. Mapping the presence of the Tad pili clusters in a phylogenomic tree made using FastANI and a selection of the most diverse species of the *Bradyrhizobium* branch (Supplementary Table S3) shows that almost all species analyzed have a complete copy of clusters 1 and 2, being the clusters 3 and 4 scattered among the different species, with cluster 4 more present in the *B. diazoefficiens* and *B. japonicum* clades (Supple-mentary Fig. S3 and Table S3). This result suggests that if clusters 1 and 2 originated from a duplication event, it probably occurred in a common ancestor of *Bradyrhizobium*. This finding is consistent with the high degree of conservation and synteny observed in previous analysis (Supplementary Figs. S1 to S3).

### 2. Tad pilus affects biofilm formation in *B. diazoefficiens* USDA 110

Wide conservation of these clusters suggested physiological and ecological importance of pili in *Bradyrhizobium*. To unravel the role of pili in *B. diazoefficiens* USDA 110, we generated deletional mutants of pili clusters 1, 2 and 3 (TP1, TP2 and TP3)(Supplementary Fig. S4). Cluster 4 (TP4) was not mutagenized because it was already studied by us (29). The mutants were tested for biofilm formation capacity, measured as adhesion in PVC 96-well microtiter plates (Fig. 2A-B). The Δpili2 mutant showed 12 to 20-fold increase in biofilm formation compared to the wild type (WT). This was unexpected, as pili are usually associated with adhesion to surfaces. In most of the bacterial models studied to date, lack of Tad pili leads to impairment of adhesion and biofilm formation (26, 32–3)). Besides, the other two mutants, namely Δpili1 and Δpili3, exhibited a behavior similar to the WT (Fig. 2A-B). These findings were confirmed by testing several clones obtained independently from different merodiploids and, in all cases, identical results were obtained (data not shown). Similar phenotype of increased biofilm formation was confirmed by direct observation of 7 day-biofilms formed on a glass slide under light microscopy after crystal violet staining (Fig. 2C).

**Figure 2.**
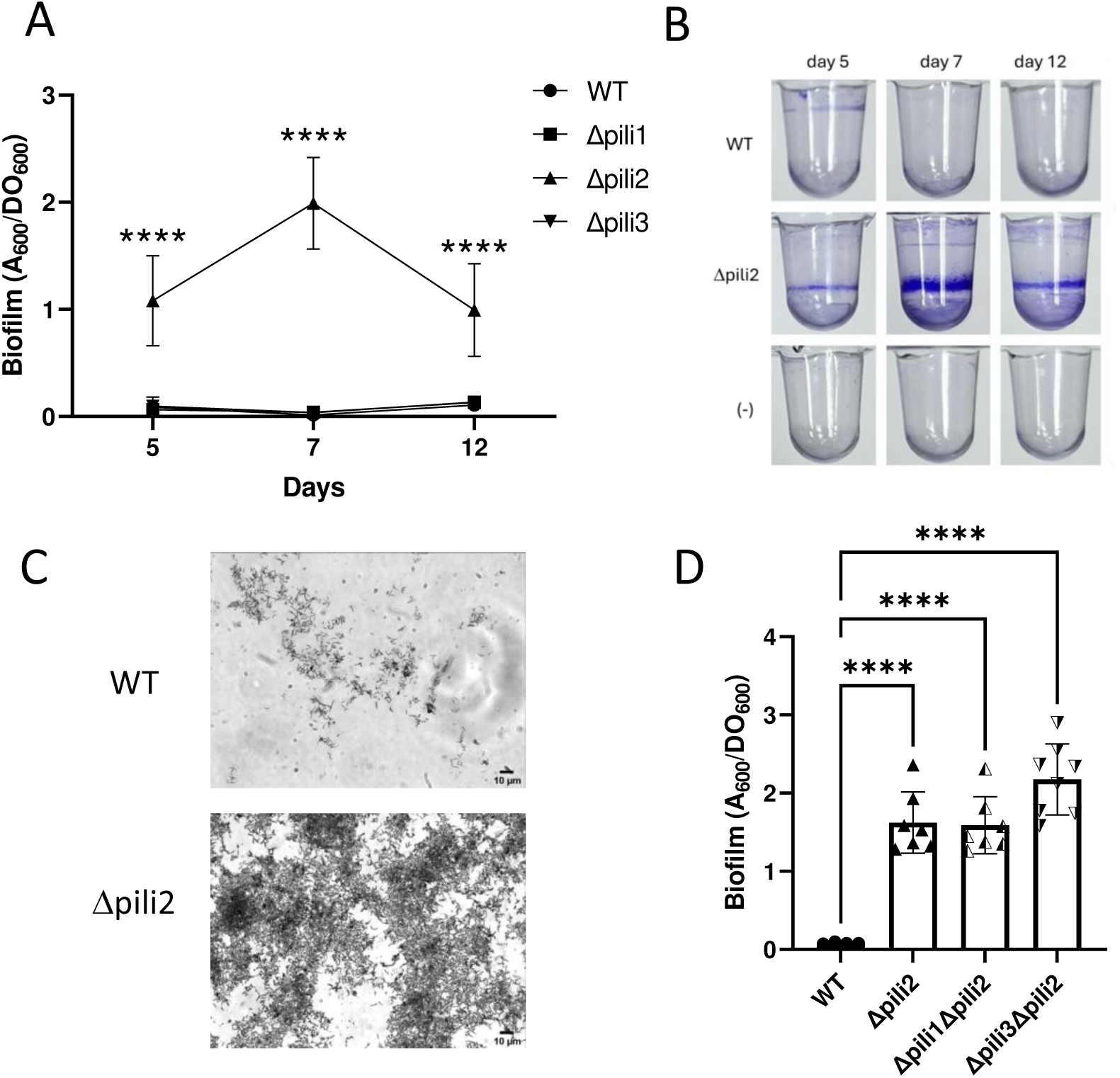
Biofilm formation by WT and Dpili mutants. **A.** Evaluation of biofilm formation at different incubation times. Data represent the results from 2 independent experiments, each with four biological replicates. **B.** Representative images of wells stained with crystal violet for the WT and the Dpili2 mutant before quantification. **C.** WT and Dpili2 mutant observed by light microscopy after crystal violet staining. **D.** Biofilm formation evaluated for the double mutants Δpili1Δpili2 and Δpili3Δpili2 compared to the WT and the Δpili2 mutant at 7 days incubation time. Statistical significance was determined by analysis of variance and Tukeýs post−hoc test comparison for differences between strains indicated. ****, indicate P <0.0001; only statistically significant differences have been indicated.

To explain this observation, we hypothesized that the augmented biofilm formation capacity in the Δpili2 mutant may be the result of deregulation or upregulation of TP3 when TP2 loses its function (39). To test this, we evaluated the biofilm formed by double mutants where TP1 and 2 (Δpili1Δpili2) or TP2 and 3 (Δpili2Δpili3) were genetically removed. Interestingly, and similarly to the single Δpili2 mutant, both double mutants increased their biofilm forming capacity (Fig. 2D). Therefore, this result excludes the possibility that the other clusters were involved in the phenotype observed in the Δpili2 mutant and reinforces the results obtained with this strain. Furthermore, the findings that i) deletion of TP1 and TP3 did not produce any effect on biofilm formation, ii) their inactivation in the Δpili2 mutant did not reverse the effect of TP2 deletion, and iii) the lack of biofilm inhibition in the Δpili2 mutant, indicate that none of the three pili are direct determinants in biofilm formation in this experimental setinng and suggest that each may have an alternative function.

In addition, the biofilms formed by fluorescent derivatives of the WT and Δpili2 strains were evaluated by confocal scanning laser microscopy (CLSM) at day 7. Overall, the biofilm architecture formed by these strains on chambered coverglass appeared slightly different (Fig. 3A). However, XZ projections showed that the WT developed a dense biofilm structure unlike the Δpili2 mutant, which formed loosely packed macro-aggregates, resulting in a biofilm crossed by multiple pores and channels. When bjGFP-WT (green) and mCherry-Δpili2 (red) were allowed to form a mixed biofilm at 1:1 ratio (Fig. 3B) the architecture of each structure was modified in comparison to the individual biofilms. Interestingly, both strains segregated in the mixed biofilm almost as frequently observed when mixing different species, forming separate clusters that contain either WT or Δpili2 mutant cells. This observation underscores the importance of Tad pili in recognition and establishment of tight cell-cell interactions. It is important to note that the rates of biofilm formation by bacteria expressing any of the fluorescent labels (bjGFP or mCherry) were similar (not shown), and no difference in biofilm formation was detected by non-fluorescence microscopy of biofilms formed by fluorescently labelled or unlabeled bacteria.

**Figure 3.**
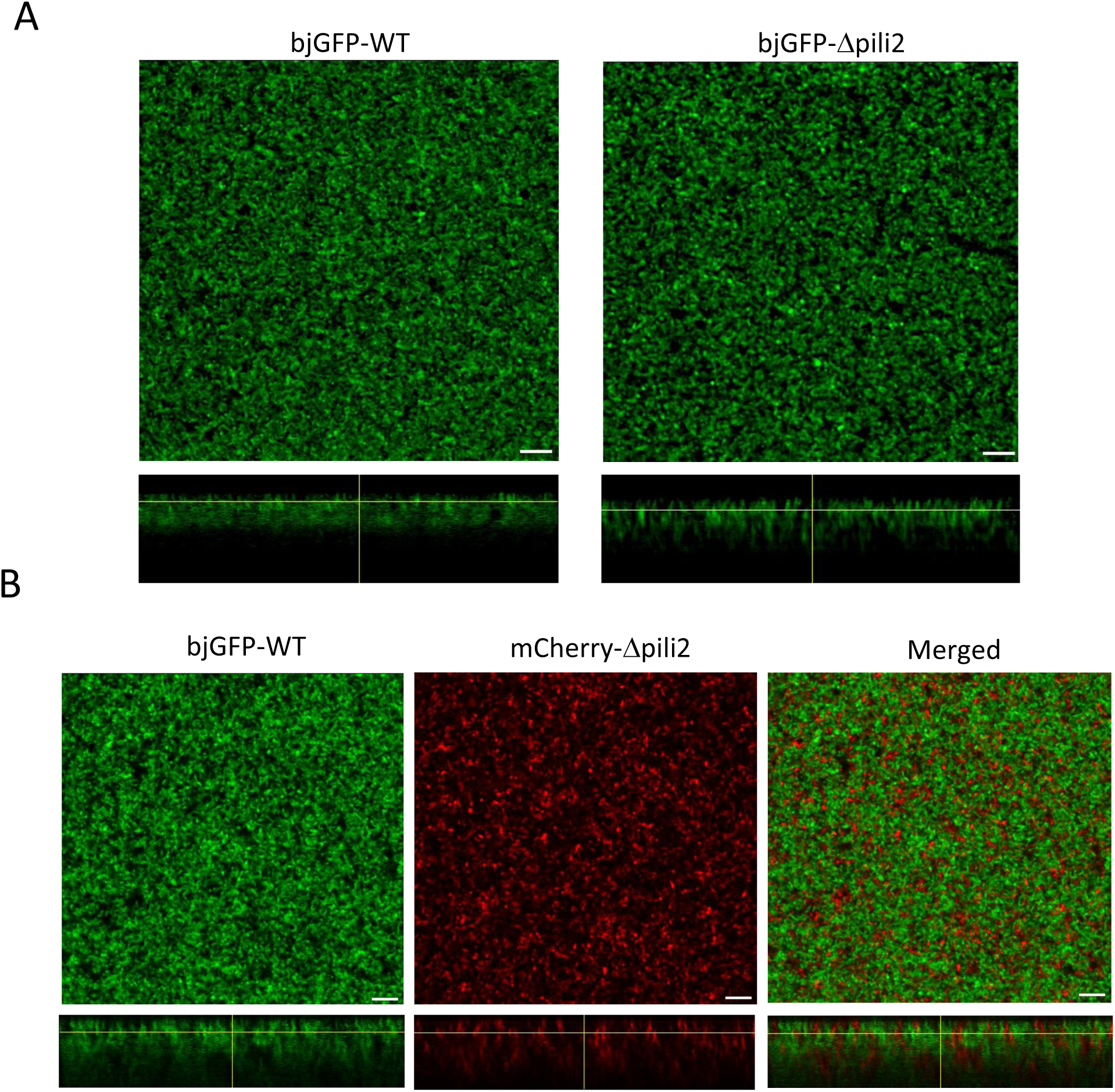
Biofilm structure analysis. CSLM images of horizontal (XY) and vertical (XZ) projections (large and bottom panels, respectively) of 7 day−biofilms formed by WT and the Δpili2 mutant. **A.** Individual biofilms formed by bjGFP−derived WT and Δpili2 strains. **B.** Mixed biofilm formed by bjGFP−WT (green) and mCherry−Δpili2 mutant (red). No interactions between the WT and mutant were observed inside the cell clusters (merged image). The glass surface is in the upper side of the xz projections; scale bars represent 10 μm. Each assay was repeated with at least two biological replicates.

### 3. Lack of Tad pili2 impairs bacterial motility

The cellular program leading to biofilm is negatively correlated with bacterial motility. In view of the different biofilm phenotypes observed above, here we evaluated the coordination of biofilm formation and motility in the context of the Tad pili mutants. First, we evaluated the swimming capacity of the mutants in semisolid medium (agar 0.3%). The Δpili2 mutant showed a substantial reduction in its swimming capacity when compared to the WT, Δpili1 and Δpili3 mutants (Fig. 4A-B). To determine if some flagellar system was affected by the pilus absence, we purified flagellins from liquid cultures and analyzed them by SDS-PAGE (Supplementary Fig. S5). This analysis showed the presence of both flagellins, and therefore this feature does not explain itself the reduced motility of Δpili2. Hence, the difference between WT and Δpili2 might reside in defective functioning of the flagellar motor or propensity to filament detachment in the Δpili2 mutant.

**Figure 4.**
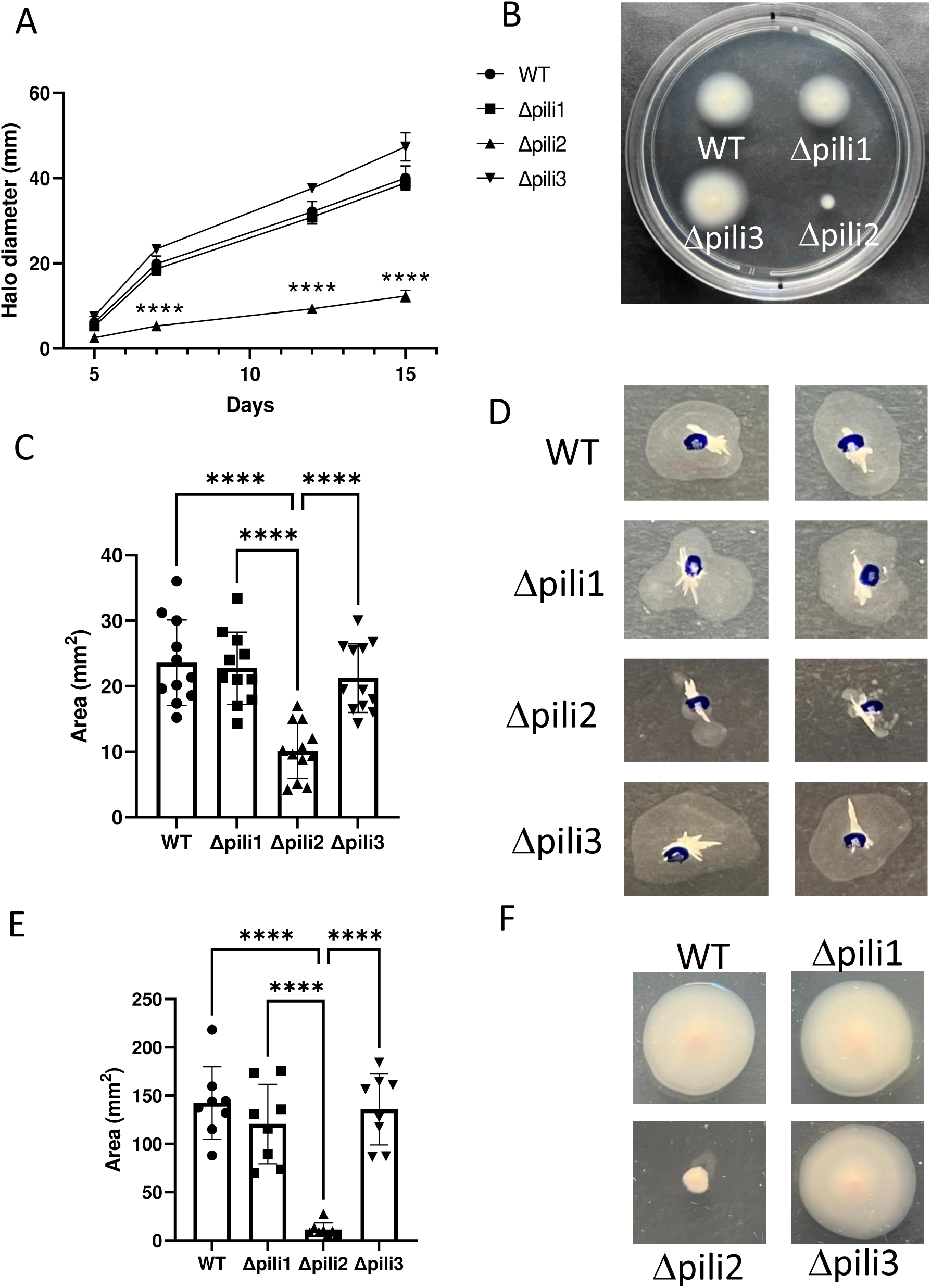
Motility characterization of the pili mutants. **A.** Swimming in semisolid AG medium (agar 0.3%) over time. **B.** Swimming plate at 7 days incubation with the WT and pili mutants. **C.** Area covered by twitching motility in the interface between the growth medium and the plate. **D.** Two representative images for each strain from the twitching experiment. **E.** Area covered by spreading motility over the surface of the growth medium. **F.** One representative image for each strain of the experiment of gliding/spreading motility. Data represent results from at least 2 independent experiments, each with minimum of four biological replicates. Statistical significance was determined by analysis of variance and Tukeýs post−hoc test comparisons for differences between strains. ****, P <0.0001; only statistically significant differences were indicated.

T4P are frequently associated with twitching motility (10, 14, 40). Specifically, in *Liberibacter crescens*, the Tad pilus has been linked to this type of movement (41), although in many other bacteria it is not responsible for this function. Given this, we evaluated this trait in *B. diazoeffi-ciens* pili mutants. As with the findings observed in the swimming experiments, only the Δpili2 mutant showed a deficiency in twitching motility in comparison to the WT (Fig. 4C-D). In addition, we observed that the Δpili2 mutant was unable to move or spread over the surface of the agar medium, (Fig. 4E-F) showing that lack of TP2 also affected the spreading motility of the mutant. The motility impairment was also confirmed for the double mutants Δpili1Δpili2 and Δpili2Δpili3 described before (data not shown). The results of this study demonstrate that the absence of pili encoded in cluster 2 in *B. diazoefficiens* USDA 110 impacts all the mechanisms of motility normally displayed by this strain, including twitching, spreading, and swimming.

### 4. Effect of carbohydrates on the Δpili2 mutant phenotype

During the analysis of biofilm formation, we observed a peculiar behavior of the Δpili2 mutant that could be linked to its phenotype of increased adhesion to PVC. Unlike the WT strain, the Δpili2 mutant does not sediment at the bottom of the well. Fig. 5A shows WT, Δpili1 and Δpili3 strains forming compact bacterial sediments at the bottom of the well after 7 days of incubation. However, this effect was not observed in the Δpili2 mutant.

**Figure 5.**
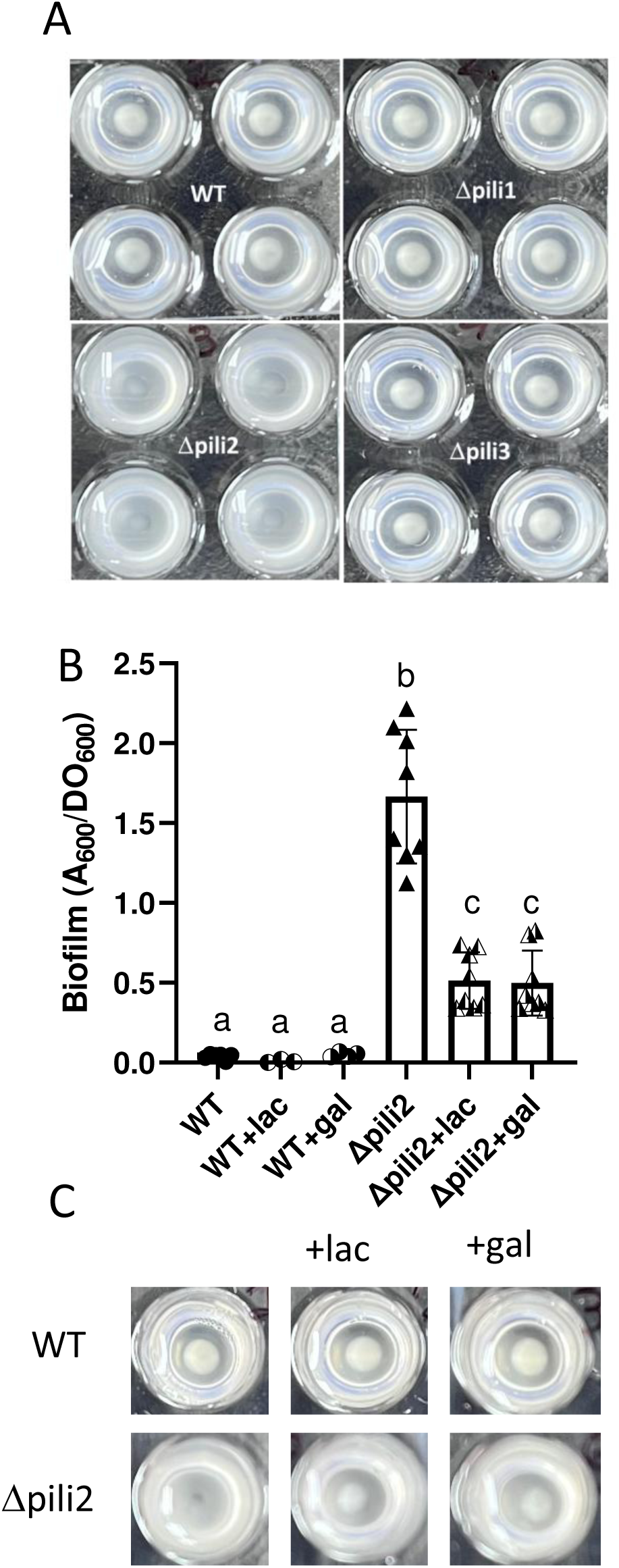
Biofilm formation inhibition assay. **A.** Images of wells from the biofilm assay in standard conditions for the WT and pili mutants before the staining treatment at 7 days of incubation. **B.** Biofilm assay for the WT and the Δpili2 mutant in presence of lactose (lac) or galactose (gal). **C.** Representative wells for the WT and the Δpili2 mutant in presence of lactose or galactose before the staining treatment after 7 days of incubation. Data represent results from 2 independent experiments, each with four biological replicates. Statisitcal significance was determined by one way ANOVA and Tukeýs post−hoc test comparison for differences between strains indicated. Different letters indicate significant differences with p <0.0001.

To investigate this phenomenon further, we evaluated biofilm formation in the presence of two different carbohydrates, lactose and galactose, which were previously described as inhibitors of bacteria-bacteria interaction mediated by TadG, TadE and/or TadF in *B. diazoefficiens* (29, 42–44). We observed a partial reversal of biofilm formed by the Δpili2 mutant in the presence of these carbohydrates (Fig. 5B). In addition, sedimentation of Δpili2 mutant cells was observed in microtiter plates in the presence of lactose and galactose (Fig. 5C). A copy of *tadGE* may be found in cluster 3 and additionally, a copy of *tadGEF* lies at cluster 4 (Fig. 1B). This last copy, from which we previously observed adhesin activity to plant roots (31), remained intact in all the mutants investigated here. Hence, we speculated that the pilus encoded in cluster 2 might in some way inhibit TadG/TadE action. In addition, cell-cell contacts mediated by these adhesins in the Δpili2 mutant might explain the exclusion of WT cells from the clusters observed in Fig. 3B.

### 5. Lack of Tad Pili does not modify EPS production

The quantity (or quality) of exopolysaccharides (EPS) produced may affect bacterial surface cell-cell interactions (45) and biofilm formation. To establish possible links between EPS production and the presence of each set of pili loci, we measured the amounts of EPS produced by all pili mutants compared to the WT strain. Fig. 6 shows the results of four independent experiments where no significant differences between the mutant strains and the WT were ob-served. Therefore, Δpili2 phenotype cannot be attributed to alteration of EPS amounts produced by these strains.

**Figure 6.**
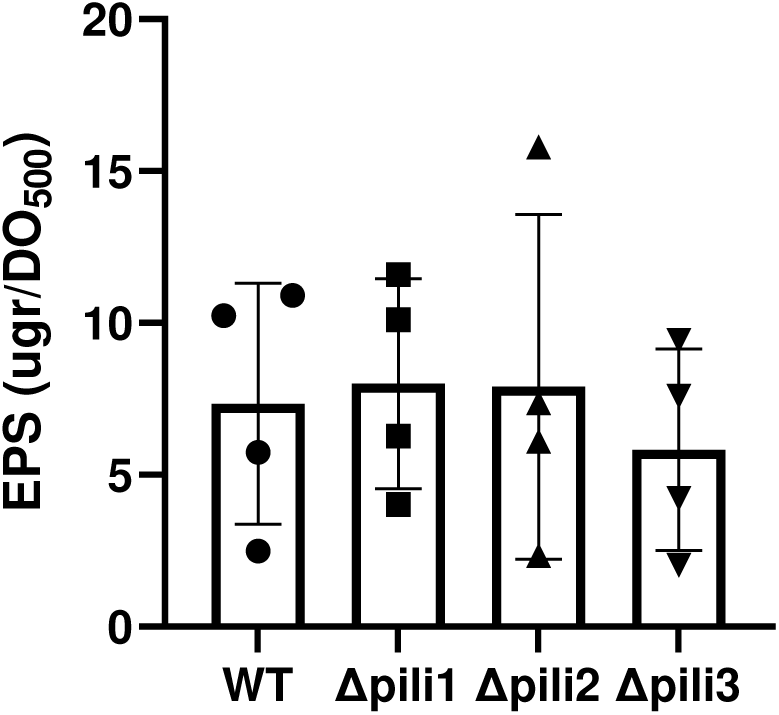
Exopolysaccharide (EPS) quantification in WT and Δpili mutants. The amounts of EPS are expressed based on total culture growth (OD_500_). Data represent results from 4 independent experiments, each with three biological replicates. EPS values are given as means ± SD. No significant differences were found by ANOVA and Tukey’s post hoc test for comparisons between strains.

### 6. The Δpili2 mutant have augmented c-di-GMP level

Up to this point, the phenotype observed in the Δpili2 mutant indicates an increase in its biofilm forming capacity. In general, the biofilm state is associated with elevation of the intracellular concentration of the c-di-GMP second messenger pool (46–4)). To test this relationship, we measured c-di-GMP levels in the WT and the Δpili mutants grown in liquid culture. As shown in Fig. 7, the intracellular level of c-di-GMP in the Δpili2 mutant was 36% higher than the WT, which suggests that the biofilm effects of this deletion might be mediated by second messen-ger levels.

**Figure 7.**
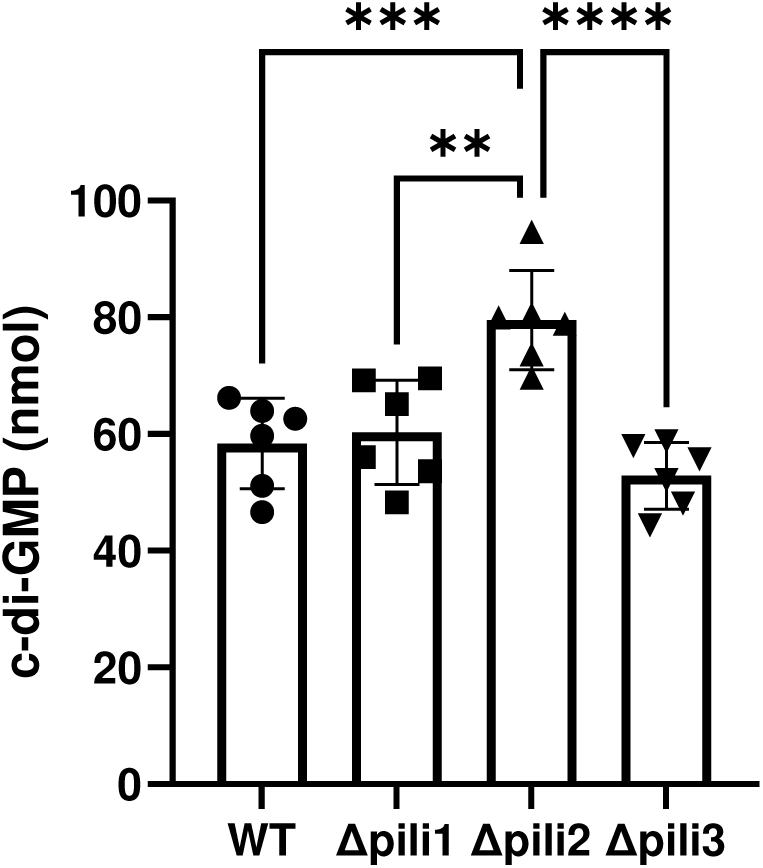
C-di-GMP quantification by HPLC-Ms/Ms identification. Data represent results from 2 independent experiments, each with three technical replicates. Statistical significance was determined by analysis of variance and Tukeýs post hoc test comparison for differences between strains indicated. **, *** and ****, indicate p<0.01, p<0.001 and p<0.0001 respectively; only statistically significant differences have been indicated.

### 7. The Flp1 pilin subunit encoded in cluster 2 is partially required for biofilm formation by Δpili2 mutant

Despite deletion of most of cluster 2, the prepilin gene *flp1* encoded in ORF *bsl1442* remained intact in the Δpili2 mutant (Supplementary Fig. S4) and therefore, it might be responsible, at least in part, for the phenotypes observed before. To address this question, we obtained a *bsl1442* deletion mutant leaving intact the remaining of cluster 2 (Δ1442 mutant strain, Supplementary Fig. S6). This mutant showed a phenotype similar to WT (Fig. 8A). It is likely that one of the paralogous genes encoding other pilins can compensate for its function in the Δ1442 mutant and could lead to a functional pili system, as observed previously in *A. tumefaciens* (26). To corroborate this statement, we obtained double mutants Δ1442 on the genetic background of the Δpili2 mutant (Δ1442Δpili2) and reciprocally, Δpili2 mutant on the genetic background of the Δ1442 mutant (Δpili2Δ1442) (both equivalent but obtained over different genetic backgrounds). These mutants are expected to not have a functional TP2 apparatus, and at the same time they should lack the Flp1 prepilin encoded in *bsl1442*. The behavior of both double mutants was intermediate between the WT and the Δpili2 mutant (Fig. 8B) indicating that, indeed, the Bsl1442 paralog has some role in biofilm formation by the Δpili2 mu-tant.

**Figure 8.**
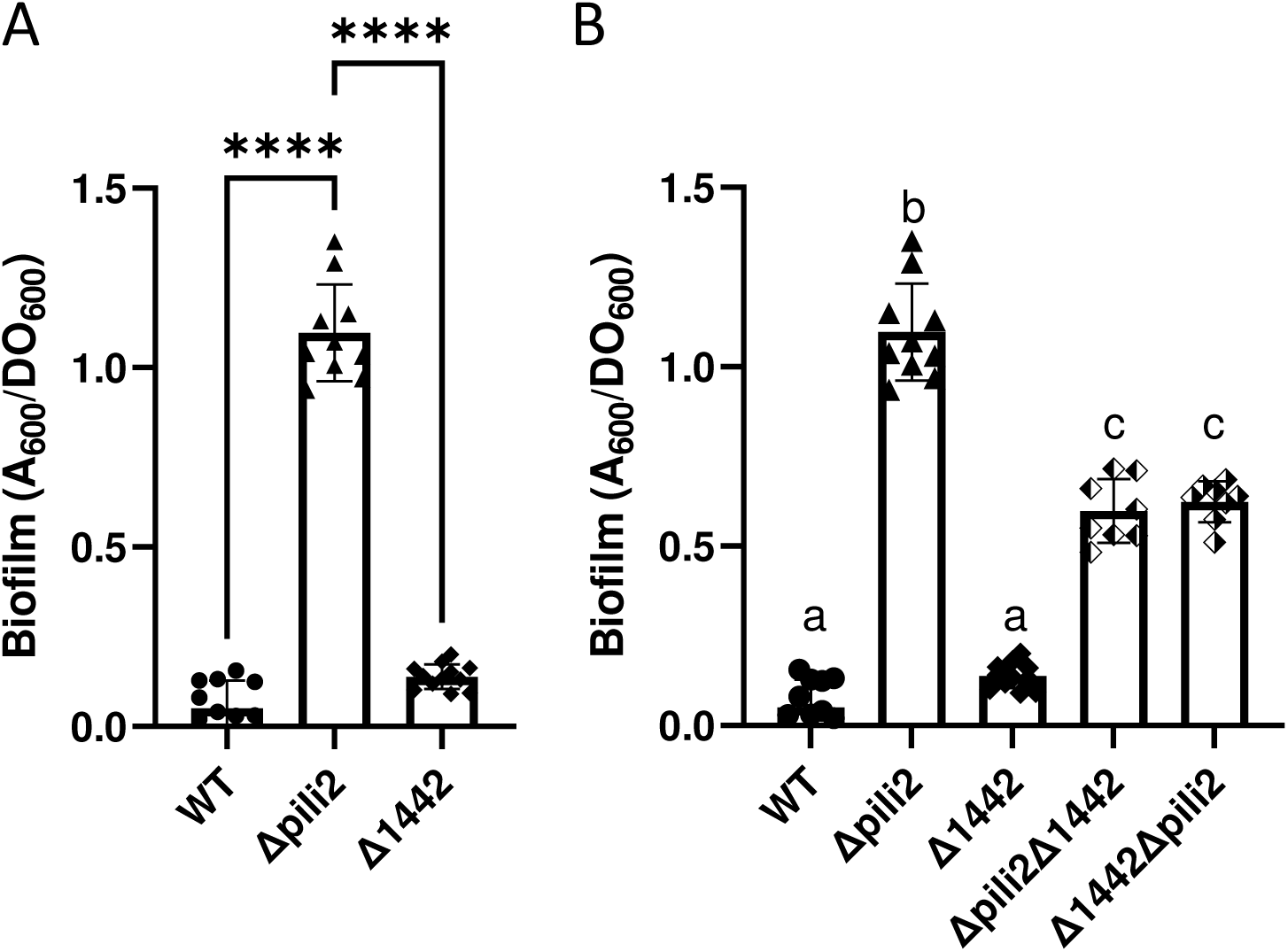
Biofilm formation in Δ1442 mutants. **A.** Biofilm formation capacity of WT and Δpili mutants measured after 7 days of incubation. **B.** Biofilm formation capacity of WT and the Δpili double mutants measured after 7 days of incubation. Data represent results from 2 independent experiments, each with four biological replicates. Statistical significance was determined by analysis of variance and Tukeýs post hoc test comparison for differences between strains indicated. **** or different letters, indicate p<0.0001; only statistically significant differences have been indicated.

### 8. The lack of tad pili does not affect soybean nodulation

One of the most interesting facts of bradyrhizobia is their capacity to symbiotically interact with host plants. At early interaction of *B. diazoefficiens* with host roots, it was observed that flagellar swimming motility is not required unless the rooting substrate is water-saturated (49) and swarming motility has no role in soil (50). In turn, despite adhesion to roots is an obvious requirement for infection and nodulation, enhancing adhesion rendered more competitive and infective bradyrhizobia (51, 52). The pili could be involved in this part of the life cycle of this microorganism at many levels. Therefore, we performed soybean nodulation experiments to determine if pili function is necessary for successful symbiosis. In plants inoculated with the WT or the Δpili2 mutant, no differences were observed in shoot dry weight, leaves chlorophyll content, numbers of nodules formed or nodules dry weight (Fig. 9A-C), thus indicating that TP2 is not required for nodulation of soybean plants.

**Figure 9.**
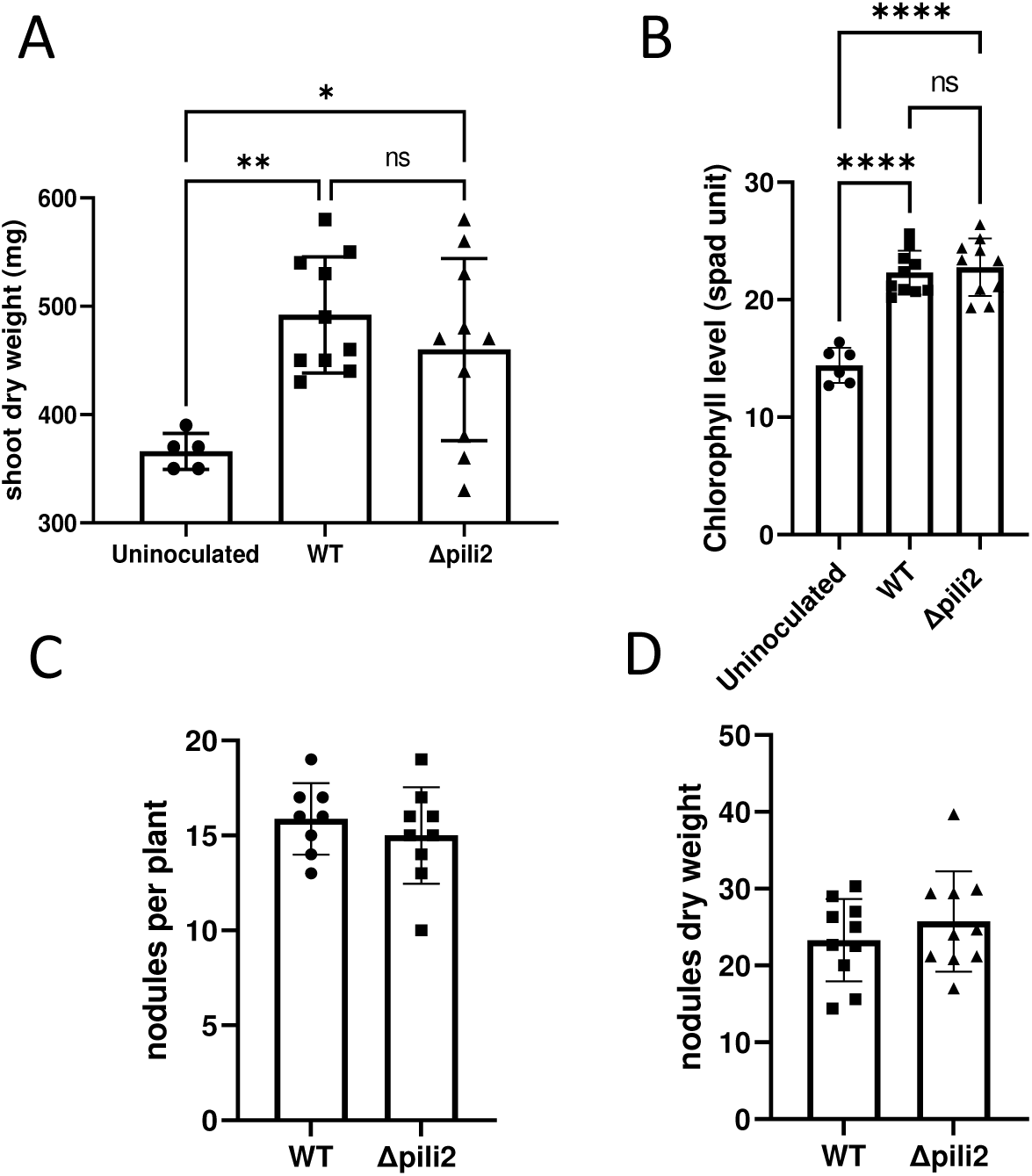
Pili mutant displays a normal symbiotic interaction with soybean. **A.** Shoot dry weight per plant. **B.** Chlorophyll content in leaves. **C.** Number of nodules per plant. **D.** Nodules dry weight per plant. Results correspond to one representative experiment from two independent repetitions, and all results are given as means ± SD. *, ** and ****, indicate p<0.05, p<0.01 and p<0.0001 respectively.

## Discussion

This work aimed at contributing to the characterization of the Tad pili apparatus in *B. diazoeffi-ciens* USDA 110, which could be extended to other species in the *Bradyrhizobium* genus, contributing to the knowledge of its role in surface sensing, adhesion, and biofilm formation. Our findings highlight the significance of Tad pili in the lifestyle of *B. diazoefficiens*, shedding light on its multifaceted role in bacterial physiology and potential applications in agriculture and biotechnology.

Our bioinformatic analyses revealed that Tad pili are widely present and highly conserved within the *Bradyrhizobium* genus (Supplementary Fig. S2), with multiple complete and partial gene clusters encoding these appendages. The high conservation, especially of clusters 1 and 2, suggests that these pili could play a critical role in the bacterial environmental adaptability, but no study was ever done in this genus. Although cluster 1 seems not complete, it is present in all but one of the strains analyzed. In addition, cluster 2, which seems complete, is also widely spread, since only three strains lack it. Cluster 3 seems also complete, but contrarily to cluster 2, it is present in roughly half of the genomes analyzed. Strikingly, there is only one example where cluster 3 is present and cluster 2 is absent, pointing to a more fundamental or ancient role of cluster 2. As for cluster 4, which is incomplete, it is the least represented.

Contrary to the phenotypes observed in other bacterial models, where the lack of pili typically reduces adhesion and biofilm formation, the Δpili2 mutant in *B. diazoefficiens* USDA 110 exhib-ited an increased biofilm formation capacity. This unexpected result suggests that the loss of functional pili (cluster 2) triggers a compensatory program leading to augmented biofilm for-mation, possibly through deregulation or upregulation of other adhesive components.

When we produced the Δpili2 mutant, we removed most of the genes encoded in cluster 2, but left intact *bsl1442*, annotated as *flp1* pilin, at the 3’ end of the cluster. This gene has its own promoter and was observed as expressed in liquid culture (53). In *C. crescentus*, one of the mechanisms that operates in the surface-sensing process involves the Tad pili and c-di-GMP signaling (24). In this way, the amount of prepilin present in the plasma membrane ap-pears to be one of the signals that activates the PleC histidine kinase pathway, generating an increase in the amount of intracellular c-di-GMP through the activation of the diguanylate cyclase, PleD. (54). This is in agreement with the results obtained by Snyder *et al.* (55) who observed that the obstruction of the pili filament motion leads to an increase in the intracellular pool of c-di-GMP, along with premature replication initiation and an increase in stalk production. These observations lead to one of the accepted models in *C. crescentus* that postulates the function of Tad pili as a mechanical surface sensor (24). This model proposes that, when the movement of the filament is blocked because it is attached to a surface, accumulation of prepilin subunits in the plasma membrane takes place (21). This accumulation works as the signal that activates the synthesis of c-di-GMP and starts the cellular adhesion program (55). In the Δpili2 mutant, the prepilin Flp1 (Bsl1442) was active, since biofilm capacity of the double mutant Δpili2Δbsl1442 diminished substantially with respect to that of the single Δpili2 mutant. Thus, it may be hypothesized that, given the lack of the pilus apparatus from cluster 2, Flp1 (Bsl1442) oversupply might accumulate, leading to the observed increase in c-di-GMP intracellular concentration. This might enhance biofilm formation, as seen in the microtiter plate, by means of another adhesin that is stimulated by the increased levels of c-di-GMP. Even if cluster 3 is active, a similar phenotype in the Δpili3 mutant was not observed, perhaps due to the interruption of the *flp1* paralogous blr3490 in this strain.

A striking effect of Δpili2 mutation was the production of large bacterial networks, which remained adhered to glass slides after washing (Fig. 2C). In parallel, this mutant did not sediment after 7 days incubation in microtiter plates, by difference with the WT or the Δpili1 or Δpili3 mutants (Fig. 5A). Similar results were obtained in *Bordetella holmesii*, where a mutant Δ*bipA* strain, which sedimented in an autoaggregation assay, was defective in biofilm formation and did not produce bacterial networks at the air-liquid interface, contrarily to the WT (56). At early steps of development, biofilms require the formation of bacterial networks, which may be formed in the liquid before settling on the surface, or after individual cells are adhered and recruit additional cells (57). Therefore, our results suggest that TP2 mediates cell-cell adhesion, which forms bacterial aggregates in *B. diazoefficiens*, preventing the formation of networks required for biofilms, which are mediated by a different adhesin. The formation of networks that prevented sedimentation of the Δpili2 mutant was partially reversed by the presence of lactose or galactose (Fig. 5C), which were previously characterized as haptens of the BJ3) bac-terial lectin (42–44). Although we could not find the complete DNA sequence proposed for this lectin in the *B. diazoefficiens* USDA 110 genome, we found part of it in the *tadG* (blr3941) ORF at cluster 4 and observed that mutating it diminished adhesion to soybean roots (29). Moreover, Ho and coworkers (42–44) described BJ3) mediates cell-cell adhesion and that this is impaired by lactose and D-galactose.

Taken together, all these results and those informed in the present paper, point to a complex interaction among pili functions in *B. diazoefficiens*, where at least clusters 2 and 4 might mediate different cell-cell contacts: while cluster 2 might be required for cell-to-cell autoaggregation, cluster 4 might mediate cell-to-cell loose contacts leading to network formation and more efficient biofilm development. This hypothesis agrees with the CLSM images in Fig. 3, where it is evident that the WT formed more compact structures than the Δpili2 mutant, and in mixed inocula, both strains were mutually excluded, perhaps because of inability of Δpili2 mutant to recognize some target in the cell surface for autoaggregation. This intricate interaction amongst pili clusters might be related to the complexity of the free-living state of this bacterium, characterized by the need to settle on surfaces as soil particles and organic remnants for survival in the soil, or on root surfaces to initiate infections. Clearly, the numbers of bacterial cells and their contacts as well as the structure of microcolonies formed should have evolved differently for adapting to the different processes that they trigger.

While the pilus apparatus should not be directly involved in the swimming and spreading motilities it could exert some kind of direct or indirect control in *B. diazoefficiens* USDA 110. The impaired motility of the Δpili2 mutant, is in agreement with its augmented biofilm formation capacity. Both characteristics could be explained by the increased level in the intracellular c-di-GMP-pool in the mutant.

Notably, soybean nodulation and N_2_ fixation were not affected by TP2 deletion (Figure 9). This suggests a more significant function of this system in free-living bacteria, however more specific parameters such as competitiveness for nodulation or infectivity will need to be revised in the future.

In conclusion, our study provides valuable insights into the genetic and functional landscape of Tad pili in *B. diazoefficiens*. The diverse roles and complex interactions of these pili in cell-to-cell autoaggregation, network formation and biofilm development highlight their importance in bacterial survival and adaptation to symbiosis. Understanding these mechanisms opens avenues for potential applications in enhancing bacterial symbiosis with plants, improving agricultural productivity, and developing novel strategies for biofilm management in industrial setinngs. Future research should focus on elucidating the specific regulatory pathways and environmental signals that modulate Tad pili functions, further enhancing our understanding of bacterial physiology and its applications.

## Material and Methods

### Bacterial strains and culture conditions

The bacterial strains and plasmids used in this work are summarized in Table S4. All *Bradyrhizobium diazoefficiens* strains were grown routinely in solid yeast extract-mannitol (YEM) (58) at 28°C. For all the assays, the strains were grown in arabinose-gluconate-supplemented HM salts (AG medium) at 28°C and 180 rpm (59). For conjugation assays, bradyrhizobia were grown in peptone-salt-yeast extract (60). *Escherichia coli* strains were grown in Luria-Bertani (LB) medium (61) at 37°C. Antibiotics were added at the following concentrations: 150 and 25 mg.ml^-1^ kanamycin (Km), 100 and 10 mg.ml^-1^ gentamicin (Gm), and 100 and 10 mg.ml^-1^ tetracycline (Tc) for *B. diazoefficiens* and *E. coli*, respectively; 200 mg.ml^-1^ ampicillin (Ap) for E. coli; and 20 mg.ml-1 chloramphenicol (Cm) for *B. diazoeffi-ciens*.

### Cloning procedures

All cloning procedures, including DNA isolation, digestion, ligation, and strain transformation, were carried out according to Sambrook et al. (2001)(61). To generate the Tad pili deletions in *B. diazoefficiens* USDA 110 were used a markerless strategy based on the pK18*mobSacB* plasmid (62). The vectors needed for mutant constructions were obtained by cloning an upstream and downstream fragment of DNA of the region planned to be deleted as indicated in Figure S5. The exact position of PCR fragment cloned, and restriction enzyme used for this porpoise are detailed in Figure S5. The PCRs reactions were carried out with Taq or Pfu polymerase (Productos Bio-Lógicos, Quilmes, Argentina) and the primers used are detailed in Table S5. The PCR fragments were agarose gel purified by an extraction kit (DSBIO^TM^) and digested with the necessary restriction enzymes (Promega^TM^) according to the supplier specifications. The pK18mob*sacB*::Δpili1, pK18mob*sacB*::Δpili2 and pK18mob*sacB*::Δpili3 plasmid (Tabla S5) obtained were transconjugated from *E. coli* S17-1 to *B. diazoefficiens* USDA 110 as was described before (62). Merodiploids of both recombinational events (upstream and downstream homologous fragments) for each pili cluster were selected by kanamycin resistance and corroborated by PCR. To obtain the deletion mutant, excision of the integrated plasmid from the merodiploid (Km^R^, Suc^S^) clones was facilitated by culturing the recombinants in AG medium without antibioitic for 24 to 48 h and then plating on YME agar supplemented with sucrose 12,5% (P/V). The resulting clones (sensible to kanamycin and resistant to sucrose) were PCR checked for the deletion event. The construction of the strains with multiple deletions was done following the same procedure but applied over the desire genetic backgrounds. To obtain the strains labelled with bjGFP or mCherry the plasmids design by Lederman et al. (2006)(63) were used.

### **Biofilm assay** (adhesion to Polyvinyl Chloride (PVC) 96-well plate)

Static biofilm formation was determined macroscopically by a quantitative assay with 96-well microtiter dishes, whereby biofilms were stained with crystal violet (CV) based on a method described previously by O’Toole and Kolter (199))(40), with minor modifications. The bacteria were grown in 10 mL AG medium, incubated with agitation for 48 h at 28°C. The cultures were then diluted with fresh medium to give an OD_500_ of 0.05. One 150 μL of the suspension were added to each well and incubated for different times at 28°C in humidity chamber to avoid excessive evaporation. Bacterial growth was quantified by measuring the OD_600_. Planktonic cells were gently removed, each well was washed twice with 200 ul of distilled water and dry for 15 min in incubator at 65°C. Then 200 ul CV aqueous solution (0.1%, w/v) was added, and staining proceeded for 15 min. Each CV-stained well was rinsed thoroughly and repeatedly with water and then scored for biofilm formation by the addition of 200 ul 33% acetic acid. The OD_600_ of solubilized CV (150 μL) was measured with a Micro Plate Reader (Benchmark Plus by Bio-Rad with Microplate Manager 5.2.1 software or FLUOstar OPTIMA by BMG Labtech with OPTIME 2.20R2 software). Sterile control cultures were made with AG medium in each experiment. The result of the estimate amounts of adhered biomass was calculated subtracting the blanks and normalized to the planktonic culture density (OD_600_).

### Microscopy analysis

Cells adhered to the glass slides were treated in the same way as the biofilm assay. Images were taken at 600X using the Nikon Eclipse E400 microscope equipped with a Nikon Coolpix 4500 digital camera.

### CLSM imaging of biofilms

Biofilms were formed on chambered coverglass (Nunc Lab-Tek) at 28°C in a 7 day-experiment to observe the architecture of single and dual-strain biofilms. GFP and mCherry fluorescent derivatives of WT and the Δpili2 mutant strains were used in the assays. Inocula were grown in 3 mL AG medium with agitation for 48 h at 28 °C, and then diluted with fresh medium to a final OD_500_ of 0,1. A 250 μL-volume of these suspensions were added to the microchamber wells and supplemented with 250 μL of fresh sterile AG medium to a final volume of 500 μL. Dual-strain cultures were inoculated with 250 μL of each strain at a ratio of 1:1. The microchambers were incubated statically at 28 °C in a humid sterile container for 7 days. Confocal images were acquired using a Leica TCS SP5 microscope with the 40x/1.5 oil-immersion objective, setting excitation wavelengths at 488 nm for GFP and at 594 nm for mCherry, while fluorescence emission was collected in the range 490-550 and 580-650 respectively. Optical sections were taken at various focal planes to create z-stacks with 0,25-μm intervals in the regions of interest; the number of slices was chosen individually for each imaged area to cover the whole biofilm. Image processing was done by using the open source Image J/ Fiji 6.3 software (64).

### Motility characterization

Swimming assay was done in semisolid AG plates (25 mL of 0.3% [wt/vol] agar in 90-mm plates) inoculated with sterile toothpicks from solid cultures in at least three separate plates and incubated at 28°C (49). The motility halo diameter was recorded for 2 weeks. At the same time, the strains were inoculated in the same agar plate and were photographed. Twitching and spreading assays were done in semisolid AG plates (25 ml of 0.8% [wt/vol] agar in 90-mm plates) inoculated with sterile toothpicks cultures crossing the surface until reaching the bottom of the plate from solid YMA cultures plates. To register the movements in the first place we record the spreading on the surface of the media. Then the colony was carefully removed with water and the bottom of the plate was photographed.

### Flagellin purification and analysis

For flagellin preparation, *B. diazoefficiens* was grown in AG liquid medium (30 mL) until optical density at 500 nm [OD_500_]=2 was reached. Supernatants were precipitated as described elsewhere (49), and each sample was resuspended in the same volume. Samples were analyzed by sodium dodecyl sulfate-polyacrylamide gel electrophoresis (SDS-PAGE) as described previously (65).

### EPS quantification

30 mL AG cultures were harvested after 48 h incubation at 28°C with agitation of 180 rpm, (final OD_500_ of 2). EPS was purified and quantified as previously described (45). For quantification, a colorimetric method using anthrone reagent was used (0.2% [w/v] anthrone in 96% [w/v] sulfuric acid), with glucose (1mg.ml^-1^) as standard.

### Endometabolome extraction and c-di-GMP quantification

Each strain was grown for 48 h in AG medium at 28 °C and 180 rpm. For endometabolome extraction, 5 mL of each strain were centrifuged at -10°C for 10 minutes at 5000 rpm, with 2 biological and 3 technical replicates per strain. The resulting pellets were frozen at -80°C until extraction. Cell lysis was performed using an ice-cold extraction solution containing equal volumes of methanol (LC-MS grade) and TE buffer (10 mM TRIZMA, 1 mM EDTA, pH 7), with a ratio of 500 µL of extraction solution per 5 mL of cell culture at an OD_500_ nm of 2. An equal volume of ice-cold chloroform was then added to the mixture, which was vortexed thoroughly and incubated for 2 hours in an Eppendorf ThermoMixer C shaker at 1000 rpm and 0°C in a cold room. Samples were subsequently centrifuged at -10°C for 10 minutes at maximum speed. The aqueous phase containing the metabolites was filtered using 13 mm PTFE 0.2 µm hydrophobic syringe filters (Fisherbrand). The extract was then stored at -80°C until analysis.

Quantitative determination of c-di-GMP was performed using an LC-MS/MS. The chromatographic separation was performed on an Agilent Infinity II 1290 HPLC system using a SeQuant ZIC-pHILIC column (150 × 2.1 mm, 5 μm particle size, peek coated, Merck) connected to a guard column of similar specificity (20 × 2.1 mm, 5 μm particle size, Phenomenex) a constant flow rate of 0.1 mL/min with mobile phase A with mobile phase comprised of 10 mM ammonium acetate in water, pH 9, supplemented with medronic acid to a final concentration of 5 μM (A) and 10 mM ammonium acetate in 90:10 acetonitrile to water, pH 9, supplemented with medronic acid to a final concentration of 5 μM (B) at 40° C. The injection volume was 5 µL. The mobile phase profile consisted of the following steps and linear gradients: 0 – 1 min constant at 75 % B; 1 – 6 min from 75 to 40 % B; 6 to 9 min constant at 40 % B; 9 – 9.1 min from 40 to 75 % B; 9.1 to 20 min constant at 75 % B. An Agilent 6495 ion funnel mass spectrometer was used in negative and positive ionization mode with an electrospray ionization source and the following conditions: ESI spray voltage 3000 V, nozzle voltage 1000 V, sheath gas 400° C at 11 l/min, nebulizer pressure 20 psig and drying gas 100° C at 11 l/min. The compound was identified based on its mass transition and retention time compared to a chemically pure standard. Chromatograms were integrated using MassHunter software (Agilent, Santa Clara, CA, USA). Absolute concentrations were determined based on an external Standard curve.

### Bioinformatics methods and analysis

*B. diazoefficiens* Tad pili clusters were identified using NCBI BLASTn software (66) with previously identified tad pili clusters from *S. meliloti* 2011, *A. tumefaciens* C58, *C. crescentus* N1000 and *A. actinomycetemcomitans* D7S-1 as query and *B. diazoefficiens* USDA110 genome sequence as subject. The function assignment was done by bi-directional BLASTp using the same reference organisms. The conservation of Tad pili clusters in*B. diazoefficiens* and the reference strains was analyzed with Clinker (67). The identified *B. diazoefficiens* USDA110 clusters were then used to search for tad pili clusters in other *Brady-rhizobium* species by means of custom python scripts. Briefly, *Bradyrhizobium* genomes were downloaded using download_gb_from_tsv.py script that uses the NCBI dataset software to download genomes in Genbank GBFF format by their accession numbers taken from a table passed as argument. The DNA sequences of all the downloaded genomes were extracted and merged into a unique fasta file to be used as subject for BLASTn search by means of gb_files_to_fasta.py script. Next, the blast_contigous_segments.py script was used to look for each of the tad pili clusters of *B. diazoefficiens* USDA110 –used as query– in the subject se-quence database. blast_contigous_segments.py script processes the BLASTn results merging neighbor HSPs to extend the homology regions, thus allowing for insertion and deletion of larger sequences in clusters. All the scripts used are available in github (https://github.com/maurijlozano/Bradyrhizobium_pili). Figure S2 was generated from the binary cluster presence/absence matrix using the clustermap function from the pyhton library Seaborn.

### Plant assay

DonMario 4800 soybean seeds were surface-sterilized by immersion in 96% (v/v) ethanol for 5 s and then in 20% (v/v) commercial bleach for 10 min, followed by six washes in sterile distilled water. Seeds were germinated on 1.5% (w/v) aqueous agar in the dark at 28°C. The nodulation assay was carried out in pots filled with vermiculite as support, which were initially watered with a mineral solution (Fåhraeus). After 30 days of culture in a chamber at 26°C with a 16 h photoperiod, the total number of nodules and the dry weight of the aerial part and nodules per plant were determined. The leaf chlorophyll content was measured with a SPAD-502 chlorophyllometer (Minolta, Japan) reading each leaflet of a trifoliate leaf before harvesting. Then, combined readings of each leaflet, were expressed in relative soil plant analysis development (SPAD) units as the average for each trifoliate leaf (6)).

## Acknowledgments

The authors are grateful to Silvana Stongiani, Luciana Cayuela and Florencia Lopez for laboratory technical assistance, and Claudio Mazo for assistance in plant experiments. JI, DC, JSS, OF, DB, IM are fellows of CONICET (Argentina). ML, PLA, MJA, AL, AS-B, JP-G and EM are members of the Scientific Researcher Career of CONICET (Argentina).

## References

1. Flemming, H.-C., van Hullebusch, E.D., Neu, T.R., Nielsen, P.H., Seviour, T., Stoodley, P., Wingender, J., Wuertz, S., 2023. The biofilm matrix: multitasking in a shared space. Nat Rev Microbiol 21, 70–86. 10.1038/s41579-022-00791-0

2. Sauer, K., Stoodley, P., Goeres, D.M., Hall-Stoodley, L., Burmølle, M., Stewart, P.S., Bjarnsholt, T., 2022. The biofilm life cycle: expanding the conceptual model of biofilm for-mation. Nat Rev Microbiol 20, 608–620. 10.1038/s41579-022-00767-0

3. Maier, B., 2021. How Physical Interactions Shape Bacterial Biofilms. Annu Rev Biophys 50, 401–417. 10.1146/annurev-biophys-062920-063646

4. Vandana, Das, S., 2022. Genetic regulation, biosynthesis and applications of extracellular polysaccharides of the biofilm matrix of bacteria. Carbohydr Polym 291, 119536. 10.1016/j.carbpol.2022.119536

5. Persat, A., 2017. Bacterial mechanotransduction. Curr Opin Microbiol 36, 1–6. 10.1016/j.mib.2016.12.002

6 -Laventie, B.-J., Jenal, U., 2020. Surface Sensing and Adaptation in Bacteria. Annu Rev Micro-biol 74, 735–760. 10.1146/annurev-micro-012120-063427

7. Belas, R., 2014. Biofilms, flagella, and mechanosensing of surfaces by bacteria. Trends Mi-crobiol 22, 517–527. 10.1016/j.tim.2014.05.002

8. Wong, G.C.L., Antani, J.D., Lele, P.P., Chen, J., Nan, B., Kühn, M.J., Persat, A., Bru, J.-L., Høy-land-Kroghsbo, N.M., Siryaporn, A., Conrad, J.C., Carrara, F., Yawata, Y., Stocker, R., V Brun, Y., Whitfield, G.B., Lee, C.K., de Anda, J., Schmidt, W.C., Golestanian, R., O’Toole, G.A., Floyd, K.A., Yildiz, F.H., Yang, S., Jin, F., Toyofuku, M., Eberl, L., Nomura, N., Zacharoff, L.A., El-Naggar, M.Y., Yalcin, S.E., Malvankar, N.S., Rojas-Andrade, M.D., Hochbaum, A.I., Yan, J., Stone, H.A., Wingreen, N.S., Bassler, B.L., Wu, Y., Xu, H., Drescher, K., Dunkel, J., 2021. Roadmap on emerg-ing concepts in the physical biology of bacterial biofilms: from surface sensing to community formation. Phys Biol 18. 10.1088/1478-3975/abdc0e

9. Pelicic, V., 2008. Type IV pili: e pluribus unum? Mol Microbiol 68, 827–837. 10.1111/j.1365-2958.2008.06197.x

10. McCallum, M., Burrows, L.L., Howell, P.L., 2019. The Dynamic Structures of the Type IV Pilus. Microbiol Spectr 7. 10.1128/microbiolspec.PSIB-0006-2018

11. Denise, R., Abby, S.S., Rocha, E.P.C., 2019. Diversification of the type IV filament superfam-ily into machines for adhesion, protein secretion, DNA uptake, and motility. PLoS Biol 17, e3000390. 10.1371/journal.pbio.3000390

12. Berne, C., Ellison, C.K., Ducret, A., Brun, Y.V., 2018. Bacterial adhesion at the single-cell level. Nat Rev Microbiol 16, 616–627. 10.1038/s41579-018-0057-5

13. Dubnau, D., Blokesch, M., 2019. Mechanisms of DNA Uptake by Naturally Competent Bac-teria. Annu Rev Genet 53, 217–237. 10.1146/annurev-genet-112618-043641

14. Ellison, C.K., Whitfield, G.B., Brun, Y.V., 2022. Type IV Pili: dynamic bacterial na-nomachines. FEMS Microbiol Rev 46, fuab053. 10.1093/femsre/fuab053

15. Tomich, M., Planet, P.J., Figurski, D.H., 2007. The tad locus: postcards from the widespread colonization island. Nat Rev Microbiol 5, 363–375. 10.1038/nrmicro1636

16. Mignolet, J., Panis, G., Viollier, P.H., 2018. More than a Tad: spatiotemporal control of Caulobacter pili. Curr Opin Microbiol 42, 79–86. 10.1016/j.mib.2017.10.017

17. Whitfield, G.B., Brun, Y.V., 2024. The type IVc pilus: just a Tad different. Curr Opin Micro-biol 79, 102468. 10.1016/j.mib.2024.102468

18. Kachlany, S.C., Planet, P.J., Bhattacharjee, M.K., Kollia, E., DeSalle, R., Fine, D.H., Figurski, D.H., 2000. Nonspecific adherence by Actinobacillus actinomycetemcomitans requires genes widespread in bacteria and archaea. J Bacteriol 182, 6169–6176. 10.1128/JB.182.21.6169-6176.2000

19. Kachlany, S.C., Planet, P.J., DeSalle, R., Fine, D.H., Figurski, D.H., 2001. Genes for tight ad-herence of Actinobacillus actinomycetemcomitans: from plaque to plague to pond scum. Trends Microbiol 9, 429–437. 10.1016/s0966-842x(01)02161-8

20. Skerker, J.M., Shapiro, L., 2000. Identification and cell cycle control of a novel pilus system in Caulobacter crescentus. EMBO J 19, 3223–3234. 10.1093/emboj/19.13.3223

21. Ellison, C.K., Kan, J., Dillard, R.S., Kysela, D.T., Ducret, A., Berne, C., Hampton, C.M., Ke, Z., Wright, E.R., Biais, N., Dalia, A.B., Brun, Y.V., 2017. Obstruction of pilus retraction stimulates bacterial surface sensing. Science 358, 535–538. 10.1126/science.aan5706

22. Hug, I., Deshpande, S., Sprecher, K.S., Pfohl, T., Jenal, U., 2017. Second messenger-mediated tactile response by a bacterial rotary motor. Science 358, 531–534. 10.1126/science.aan5353

23. Ellison, C.K., Rusch, D.B., Brun, Y.V., 2019. Flagellar Mutants Have Reduced Pilus Synthesis in Caulobacter crescentus. J Bacteriol 201. 10.1128/JB.00031-19

24. Sangermani, M., Hug, I., Sauter, N., Pfohl, T., Jenal, U., 2019. Tad Pili Play a Dynamic Role in Caulobacter crescentus Surface Colonization. mBio 10. 10.1128/mBio.01237-19

25. Hershey, D.M., Fiebig, A., Crosson, S., 2021. Flagellar Perturbations Activate Adhesion through Two Distinct Pathways in Caulobacter crescentus. mBio 12. 10.1128/mBio.03266-20

26. Wang, Y., Haitjema, C.H., Fuqua, C., 2014. The Ctp type IVb pilus locus of Agrobacterium tumefaciens directs formation of the common pili and contributes to reversible surface attachment. J Bacteriol 196, 2979–2988. 10.1128/JB.01670-14

27. Zatakia, H.M., Nelson, C.E., Syed, U.J., Scharf, B.E., 2014. ExpR coordinates the expression of symbiotically important, bundle-forming Flp pili with quorum sensing in Sinorhizobium meli-loti. Appl Environ Microbiol 80, 2429–2439. 10.1128/AEM.04088-13

28 -Carvia-Hermoso, C., Cuéllar, V., Bernabéu-Roda, L.M., van Dillewijn, P., Soto, M.J., 2024. Sinorhizobium meliloti GR4 Produces Chromosomal-and pSymA-Encoded Type IVc Pili That Influence the Interaction with Alfalfa Plants. Plants (Basel8 13. 10.3390/plants13050628

29. Mongiardini, E.J., Parisi, G.D., Quelas, J.I., Lodeiro, A.R., 2016. The tight-adhesion proteins TadGEF of Bradyrhizobium diazoefficiens USDA 110 are involved in cell adhesion and infectivity on soybean roots. Microbiol Res 182, 80–88. 10.1016/j.micres.2015.10.001

30. Park, S., Sauer, K., 2022. Controlling Biofilm Development Through Cyclic di-GMP Signaling. Adv Exp Med Biol 1386, 69–94. 10.1007/978-3-031-08491-1_3

31. Vesper, S.J., Bauer, W.D., 1986. Role of Pili (Fimbriae) in Attachment of Bradyrhizobium japonicum to Soybean Roots. Appl Environ Microbiol 52, 134–141. 10.112/aem.52.1.134-141.1986

32. Perez, B.A., Planet, P.J., Kachlany, S.C., Tomich, M., Fine, D.H., Figurski, D.H., 2006. Genetic analysis of the requirement for flp-2, tadV, and rcpB in Actinobacillus actinomycetemcomitans biofilm formation. J Bacteriol 188, 6361–6375. 10.1128/JB.00496-06

33. de Bentzmann, S., Aurouze, M., Ball, G., Filloux, A., 2006. FppA, a novel Pseudomonas ae-ruginosa prepilin peptidase involved in assembly of type IVb pili. J Bacteriol 188, 4851–4860. 10.1128/JB.00345-06

34. Wisniewski-Dyé, F., Borziak, K., Khalsa-Moyers, G., Alexandre, G., Sukharnikov, L.O., Wui-chet, K., Hurst, G.B., McDonald, W.H., Robertson, J.S., Barbe, V., Calteau, A., Rouy, Z., Mangenot, S., Prigent-Combaret, C., Normand, P., Boyer, M., Siguier, P., Dessaux, Y., Elmerich, C., Condemine, G., Krishnen, G., Kennedy, I., Paterson, A.H., González, V., Mavingui, P., Zhulin, I.B., 2011. Azospirillum genomes reveal transition of bacteria from aquatic to terrestrial envi-ronments. PLoS Genet 7, e1002430. 10.1371/journal.pgen.1002430

35. Pu, M., Rowe-Magnus, D.A., 2018. A Tad pilus promotes the establishment and resistance of Vibrio vulnificus biofilms to mechanical clearance. NPJ Biofilms Microbiomes 4, 10. 10.1038/s41522-018-0052-7

36. Fiebig, A., 2019. Role of Caulobacter Cell Surface Structures in Colonization of the Air-Liquid Interface. J Bacteriol 201. 10.1128/JB.00064-19

37. Andrade, M., Wang, N., 2019. The Tad Pilus Apparatus of “Candidatus Liberibacter asiati-cus” and Its Regulation by VisNR. Mol Plant Microbe Interact 32, 1175–1187. 10.1094/MPMI-02-19-0052-R

38. Tekedar, H.C., Patel, F., Blom, J., Griffin, M.J., Waldbieser, G.C., Kumru, S., Abdelhamed, H., Dharan, V., Hanson, L.A., Lawrence, M.L., 2024. Tad pili contribute to the virulence and biofilm formation of virulent Aeromonas hydrophila. Front Cell Infect Microbiol 14, 1425624. 10.3389/fcimb.2024.1425624

39. Pu, M., Duriez, P., Arazi, M., Rowe-Magnus, D.A., 2018. A conserved tad pilus promotes Vibrio vulnificus oyster colonization. Environ Microbiol 20, 828–841. 10.1111/1462-2920.14025

40. O’Toole, G.A., Kolter, R., 1998. Flagellar and twitching motility are necessary for Pseudo-monas aeruginosa biofilm development. Mol Microbiol 30, 295–304. 10.1046/j.1365-2958.1998.01062.x

41. Cai, L., Jain, M., Sena-Vélez, M., Jones, K.M., Fleites, L.A., Heck, M., Gabriel, D.W., 2021. Tad pilus-mediated twitching motility is essential for DNA uptake and survival of Liberibacters. PLoS One 16, e0258583. 10.1371/journal.pone.0258583

42. Ho, S.C., Wang, J.L., Schindler, M., 1990. Carbohydrate binding activities of Bradyrhizobium japonicum. I. Saccharide-specific inhibition of homotypic and heterotypic adhesion. J Cell Biol 111, 1631–1638. 10.1083/jcb.111.4.1631

43. Ho, S.C., Schindler, M., Wang, J.L., 1990. Carbohydrate binding activities of Bradyrhizobium japonicum. II. Isolation and characterization of a galactose-specific lectin. J Cell Biol 111, 1639– 1643. 10.1083/jcb.111.4.1639

44. Loh, J.T., Ho, S.C., de Feijter, A.W., Wang, J.L., Schindler, M., 1993. Carbohydrate binding activities of Bradyrhizobium japonicum: unipolar localization of the lectin BJ38 on the bacterial cell surface. Proc Natl Acad Sci U S A 90, 3033–3037. 10.1073/pnas.90.7.3033

45. Quelas, J.I., López-García, S.L., Casabuono, A., Althabegoiti, M.J., Mongiardini, E.J., Pérez-Giménez, J., Couto, A., Lodeiro, A.R., 2006. Effects of N-starvation and C-source on Bradyrhizo-bium japonicum exopolysaccharide production and composition, and bacterial infectivity to soybean roots. Arch Microbiol 186, 119–128. 10.1007/s00203-006-0127-3

46. Jenal, U., Reinders, A., Lori, C., 2017. Cyclic di-GMP: second messenger extraordinaire. Nat Rev Microbiol 15, 271–284. 10.1038/nrmicro.2016.190

47. Dahlstrom, K.M., Collins, A.J., Doing, G., Taroni, J.N., Gauvin, T.J., Greene, C.S., Hogan, D.A., O’Toole, G.A., 2018. A Multimodal Strategy Used by a Large c-di-GMP Network. J Bacteri-ol 200. 10.1128/JB.00703-17

48. Junkermeier, E.H., Hengge, R., 2023. Local signaling enhances output specificity of bacteri-al c-di-GMP signaling networks. Microlife 4, uqad026. 10.1093/femsml/uqad026

49. Althabegoiti MJ, Covelli JM, Pérez-Giménez J, Quelas JI, Mongiardini EJ, Lopez MF, López-Garcia SL, Lodeiro AR. 2011. Analysis of the role of the two flagella of Bradyrhizobium japoni-cum in competition for nodulation of soybean. FEMS Microbiol Lett 319:133–139. 10.1111/j.1574-6968.2011.02280.x.

50. Covelli, J.M., Althabegoiti, M.J., López, M.F., Lodeiro, A.R., 2013. Swarming motility in Bradyrhizobium japonicum. Res Microbiol 164, 136–144. 10.1016/j.resmic.2012.10.014

51. Lodeiro, A.R., López-García, S.L., Vázquez, T.E., Favelukes, G., 2000. Stimulation of adhe-siveness, infectivity, and competitiveness for nodulation of Bradyrhizobium japonicum by its pretreatment with soybean seed lectin. FEMS Microbiol Lett 188, 177–184. 10.1111/j.1574-6968.2000.tb09190.x

52. López-García, S.L., Vázquez, T.E., Favelukes, G., Lodeiro, A.R., 2001. Improved soybean root association of N-starved Bradyrhizobium japonicum. J Bacteriol 183, 7241–7252. 10.1128/JB.183.24.7241-7252.2001

53 – Čuklina, J., Hahn, J., Imakaev, M., Omasits, U., Förstner, K.U., Ljubimov, N., Goebel, M., Pessi, G., Fischer, H.-M., Ahrens, C.H., Gelfand, M.S., Evguenieva-Hackenberg, E., 2016. Ge-nome-wide transcription start site mapping of Bradyrhizobium japonicum grown free-living or in symbiosis. a rich resource to identify new transcripts, proteins and to study gene regulation. BMC Genomics 17, 302. 10.1186/s12864-016-2602-9

54. Del Medico, L., Cerletin, D., Schächle, P., Christen, M., Christen, B., 2020. The type IV pilin PilA couples surface attachment and cell-cycle initiation in Caulobacter crescentus. Proc Natl Acad Sci U S A 117, 9546–9553. 10.1073/pnas.1920143117

55. Snyder, R.A., Ellison, C.K., Severin, G.B., Whitfield, G.B., Waters, C.M., Brun, Y.V., 2020. Surface sensing stimulates cellular differentiation in Caulobacter crescentus. Proc Natl Acad Sci U S A 117, 17984–17991. 10.1073/pnas.1920291117

56. Hiramatsu, Y., Saito, M., Otsuka, N., Suzuki, E., Watanabe, M., Shibayama, K., Kamachi, K., 2016. BipA Is Associated with Preventing Autoagglutination and Promoting Biofilm Formation in Bordetella holmesii. PLoS One 11, e0159999. 10.1371/journal.pone.0159999

57. Trunk, T., Khalil, H.S., Leo, J.C., 2018. Bacterial autoaggregation. AIMS Microbiol 4, 140–164. 10.3934/microbiol.2018.1.140

58. Vincent JM. 1970. A manual for the practical study of the root nodule bacteria. IBP hand-book 15. Blackwell Scientific Publications, Oxford, United Kingdom.

59. Mengucci, F., Dardis, C., Mongiardini, E.J., Althabegoiti, M.J., Partridge, J.D., Kojima, S., Homma, M., Quelas, J.I., Lodeiro, A.R., 2020. Characterization of FliL Proteins in Bradyrhizobi-um diazoefficiens: Lateral FliL Supports Swimming Motility, and Subpolar FliL Modulates the Lateral Flagellar System. J Bacteriol 202. 10.1128/JB.00708-19

60. Regensburger, B., Hennecke, H., 1983. RNA polymerase from Rhizobium japonicum. Arch Microbiol 135, 103–109. 10.1007/BF00408017

61. Sambrook J, Russell DW. 2001. Molecular cloning: a laboratory manual, 3rd ed. Cold Spring Harbor Laboratory Press, Cold Spring Harbor, NY.

62. Dardis, C., Quelas, J.I., Mengucci, F., Althabegoiti, M.J., Lodeiro, A.R., Mongiardini, E.J., 2021. Dual Control of Flagellar Synthesis and Exopolysaccharide Production by FlbD-FliX Class II Regulatory Proteins in Bradyrhizobium diazoefficiens. J Bacteriol 203. 10.1128/JB.00403-20

63. Ledermann, R., Bartsch, I., Remus-Emsermann, M.N., Vorholt, J.A., Fischer, H.-M., 2015. Stable Fluorescent and Enzymatic Tagging of Bradyrhizobium diazoefficiens to Analyze Host-Plant Infection and Colonization. Mol Plant Microbe Interact 28, 959–967. 10.1094/MPMI-03-15-0054-TA

64. Schindelin, J., Arganda-Carreras, I., Frise, E., et al. Fiji: an open-source platform for biologi-cal-image analysis. Nat Methods 9, 676–682 (2012). 10.1038/nmeth.2019

65. Laemmli UK. 1970. Cleavage of structural proteins during the assembly of the head of bacteriophage T4. Nature 227:680–685. 10.1038/227680a0

66. Morgulis, A., Coulouris, G., Raytselis, Y., Madden, T.L., Agarwala, R., Schäffer, A.A., 2008. Database indexing for production MegaBLAST searches. Bioinformatics 24, 1757–1764. 10.1093/bioinformatics/btn322

67. Gilchrist, C.L.M., Chooi, Y.-H., 2021. clinker & clustermap.js: automatic generation of gene cluster comparison figures. Bioinformatics 37, 2473–2475. 10.1093/bioinformatics/btab007

68. Brignoli D, Frickel-Critto E, Sandobal TJ, Balda RS, Castells CB, Mongiardini EJ, Pérez-Giménez J and Lodeiro AR (2024) Quality control of Bradyrhizobium inoculant strains: detection of nosZ and correlation of symbiotic efficiency with soybean leaf chlorophyll levels. Front. Agron. 6:1336433. 10.3389/fagro.2024.1336433

